# Measuring the impact of an interdisciplinary learning project on nursing, architecture and landscape design students’ empathy

**DOI:** 10.1101/605097

**Authors:** Samantha Donnelly, Suzanne J Dean, Shohreh Razavy, Tracy Levett-Jones

## Abstract

In Australia and internationally, domestic violence is a major cause of homelessness for women and children. When designing emergency accommodation, the concerns, preferences, and perspectives of individuals who access refuge services must be sought in order to create spaces that are conducive to the needs of this diverse and vulnerable group. An empathic ‘lens’ can provide meaningful insights that can inform the design of refuge services specifically targeted at addressing these needs.

This paper describes an authentic interdisciplinary learning experience for nursing, architecture and landscape students’, and presents the results of a study designed to measure the impact of this initiative on participants’ empathy towards women and children who access refuge services as a result of homelessness and/or domestic violence. Empathy levels were measured using the Comprehensive State Empathy Scale.

The learning experience consisted of collaborative meetings with stakeholders from the refuge sector, fieldwork, individual research, exchanging ideas and problem-solving in teams. Students then developed design guides for refuges that demonstrated their emerging understanding of the specific needs and perspectives of the issues faced by women and children who were homeless.

A convenience sample of 48 students (nursing n = 22; architecture n = 11; and landscape n = 13) participated in the study. Participants were aged from 19 to 37 years with an average of 23.8 years (SD= 3.65). Pre-post Comprehensive State Empathy Scale results indicated that the empathy levels of nursing and landscape students increased and those of architecture students decreased, however, these results were not statistically significant.

In Australia, one in six women have experienced domestic violence and domestic violence remains the single largest cause of homelessness for women. Yet reports suggest that these women frequently encounter discrimination, both in the community and when accessing services. As empathy is one of the strongest negative correlates of prejudice, authentic teaching and learning activities, such as the one described in this paper, have the potential to positively impact the lived experience of these women.

## Introduction

In Australia, domestic violence is a major cause of homelessness for women and children [1]. Refuge services have been implemented to provide safe crisis accommodation, emotional support and advocacy for this vulnerable group of individuals [2]. When considering the design of refuges, an interdisciplinary and collaborative approach is needed, with input from a range of stakeholders. Healthcare professionals can provide important insights as family violence and homelessness are major causes of physical and mental harm. The contribution from architecture and landscape is also essential because of the need for good design principles to be used in creating both permanent and temporary accommodation. However, most importantly, the concerns, preferences, and perspectives of individuals who access and provide refuge services must be sought in order to create safe spaces that are conducive to the needs of this diverse group.

This paper describes an authentic interdisciplinary learning experience where nursing, landscape and architecture students engaged with refuge service providers and women who were victims of domestic violence, to gain insights into their needs and preferences in relation to the design of women’s refuges. This real-world learning experience provided an empathic ‘lens’ and enabled students to appreciate the lived experience of service users. Additionally, by working collaboratively with other disciplines, sharing knowledge, understandings and ideas, they were able to imagine possible alternatives to existing refuge spaces.

This unique learning experience was informed by the philosophical tenets of empathy. Educational interventions designed to foster empathy are increasingly being introduced into undergraduate curricula. These types of initiatives to allow learners to ‘step into another person’s shoes’ in order to gain new insights into their feelings, perspectives and experiences [3–4]. While previous studies have reported benefits of empathy in the fields of nursing [5] and more recently in the fields of architecture and design studio learning [6], limited studies have examined the impact of collaborative learning experiences involving various disciplines working together to develop real-world understandings of the lived experience of homelessness and domestic violence. The impact of this interdisciplinary learning experience on nursing, landscape and architecture students’ empathy levels were measured using the Comprehensive State Empathy Scale (CSES).

## Literature Review

The word empathy comes from the German term, Einfühlung, meaning ‘feeling into’ [7]. The theory of Einfühlung first appeared in philosophy in 1778 with Herder writing on “body truth” and was further examined by Vischer in 1873 who stated that “the result of the spatial relationship with an object is an increase or decrease in our vital sensation.” Further studies by Schmarsow and Wolfflin in 1886 explored the connection between breathing and cardiac movements with spatial experience [8]. In the early 1900s, empathy was understood as a projection of feelings onto the world. In the 1950s however, the experimental psychologist Rosalind Cartwright proposed empathy as an interpersonal exchange, rejecting projection in preference for “true” empathy – the accurate understanding of another’s feelings. More recently the discovery of mirror neurones by neuroscientists has linked empathy with personal relationships [9].

In contemporary discourse, empathy is commonly understood as the ability to accurately understand or feel what another person is experiencing, or to be able to place oneself in their context. Empathy requires the perception and articulation of the emotional state of another and results in a range of responses, often including a desire to help or an ability to experience a similar emotional state. There are two principal issues that need to be understood in the context of this research - the difference between “cognitive” and “affective” empathy, and whether empathy is understood as a “trait” or “state” condition [10].

Cognitive empathy refers to the accurate perception of another person’s perspective or mental state (feeling into), whereas affective empathy involves an ability to experience actual feelings or emotions that are consistent with another person, that is, “feeling with” or “feeling concern for” the other [10]. Krznaric proposes that empathy is the combination of the ability to perceive and share (affective) as well as to think and act (cognitive). This kind of empathetic relationship describes a balance between looking inwards and looking outwards; and stepping outside of ourselves to explore the perspectives of another person people [11].

Bloom argues, controversially, that affective empathy is often privileged over cognitive empathy, leading to inaccurate decisions, based on incorrect perceptions of another’s emotional state or value judgements of another’s experience. Bloom proposes that cognitive empathy - the ability to perceive and act from an objective position, requires understanding another, without needing to emotionally match the other [12]. This is a perspective reiterated by Zurek [13]. Empathy is also viewed as both a ‘trait’ (predisposed psychological condition or disposition) and ‘state’ (an empathetic response at a specific point in time) [14].

Empathy is an important learning outcome for future healthcare and design professionals. In healthcare, empathy is integral to therapeutic relationships, allowing nurses to respond appropriately to the emotional needs of patients, to recognize their psychological status and to respond with understanding and insight [15]. A body of research attests to the benefits of empathic interactions between healthcare professionals and patients. Such benefits include decreased levels of patient anxiety and depression [16], and increased levels of emotional wellbeing, satisfaction and adherence to treatment regimens [17}. A wide range of physiological outcomes have also been attributed to empathic encounters including improved wound healing, level of immunity, and cancer survival rates [18]; and lower blood pressure levels, pain diabetic complications [19]. Empathic interactions also enhance healthcare professionals’ job satisfaction, resilience and coping skills [16–17].

Integration of empathy in design programs has become increasingly common in recent years, although it is not often promoted as a graduate attribute, being referenced almost exclusively in relation to understanding user requirements rather than as a design skill. Focus on user-focused design thinking has shown that empathy aids in developing an understanding of the problem that can lead to more successful design outcomes including objects, buildings and urban environments. A spatial understanding in terms of empathetic response tends to involve “an attempt to trace both aesthetic effect and the meaning of spatial form … back to the human body and its sensations” [20].

In terms of landscape design, empathetic understanding involves a relationship between subject and nature. Nowak describes the human tendency to anthropomorphize and animate nature, facilitating an understanding of strange phenomena and ascribing life and feelings to the inanimate elements of nature [7]. This suggests a more constructed idea of empathetic understanding that relates to material and spatial function.

Lifchez [21], Ghajargar and Longo [22] both discuss the manifestation of empathy in the design process and the resulting communication between designer and client. Hess and Fila explore the notion that visualisation of empathetic thinking is demonstrated by students’ ability to integrate user requirements in the design brief development of a project and then in design outcomes, describing these as “empathetic pathways” [6].

In neither healthcare nor design studies, has the value of empathy within a collaborative learning experience involving nursing, landscape and architecture students working together to develop real-world understandings of the lived experience of homelessness and domestic violence been explored. Thus, the project described in this paper sought to address this gap in the literature.

## Methodology

### Aim

The aim of this study was to examine the impact of an interdisciplinary learning experience on nursing, architecture and design students’ empathy towards women seeking refuge services.

### Study design

This study employed a three-group pre-test post-test design with empathy levels measured at baseline (pre-test) and after the learning experience (post-test).

### Setting and participants

A convenience sample of nursing, architecture and design students from one Australian metropolitan university were recruited for the study.

### Ethical Consideration

Following ethics approval from the university ethics committee, potential participants were recruited using an announcement posted on an electronic learning management system (BlackboardTM). Interested students were asked to email one of the researchers if they wished to participate in the study. Although the learning activity was mandatory, only students who provided written informed consent were included the study. Data were collected between March and June 2018.

### Intervention - Learning experience

Over the semester, students undertaking the interdisciplinary learning experience were enrolled in separate nursing, architecture or landscape design subjects. The nursing students were enrolled in a third-year elective subject focused on women’s health and including content on domestic and family violence and homelessness. The subject is delivered within a feminist framework, looking closely at the rationale for gendered healthcare services for women, and in the context of a social model of health. Students undertake a four-week work-based placement in various women’s health service environments (such as women’s health clinics, sexual health clinics, abortion services, women’s substance abuse facilities such as detox and rehab centres, women’s refuges) as a part of the learning experience.

The architecture students were enrolled in a masters level elective subject that focused on the typology of refuge, exploring retreats, monasteries, sanatoriums and domestic spaces as precedents. Statistical research and current media coverage of domestic violence and refuge provision were examined with a focus on spatial provisions and programmatic requirements.

Landscape students were enrolled in a third-year design studio, which required the development of a scheme for a real-life refuge. This subject required innovative responses to community and stakeholder requirements for community landscape projects with an exploration of the biological and therapeutic characteristics of plants.

The students from all disciplines met on three occasions to collaborate about the design of the women’s refuge. The first meeting involved students working in mixed discipline teams to research the healthcare and social needs of women and children who accessed refuges, identifying typical scenarios for women accessing refuge services, and exploring their physical, mental, emotional and spatial needs. The knowledge acquired by the nursing students during their placement experiences was shared with the design students, so as to help them understand and frame user profiles in a more empathic and realistic manner. This exercise aimed to provide foundational knowledge about the varied needs of refuge users.

The second meeting took place at a local council with a service provider and case manager of the local women’s refuge. Fieldwork and conversations with clients, service providers and refuge workers allowed the real-world and current conditions of those experiencing and working in the field of domestic and family violence to emerge. The service providers also discussed the daily management and funding requirements of refuges while the case workers spoke about the lived experiences of women and children staying in the refuge.

Using the information provided by the nursing students and the insights gained through their fieldwork experiences with service providers, the architecture and landscape students then drafted spatial responses to the needs of users, with reference to safety, access to sunlight, cross-ventilation and flexible room layouts. An example of this included the design of external play areas with specific planting to address the needs of children experiencing anxiety and depression.

In the third collaborative meeting architecture and landscape students sought feedback from the nursing students, service providers and users about their spatial responses. The students showcased their sketches, plan, diagrams and design guides containing ten best-practice principles for refuges, and combining landscape and architectural elements as a type of ‘how-to guide’ for conceptualising a refuge as a designed space.

### Data collection

Students’ empathy levels were measured using the Comprehensive State Empathy Scale (CSES) [24] prior to and immediately following the learning experience. Demographic data were also collected prior to the learning intervention. The CSES was previously developed to evaluate the impact of an educational experience on participants’ empathy towards specific people or groups over a brief time period. It measures both cognitive and affective empathy, as well as state empathy (at a point in time). [25] The scale includes 30 items and takes 10-15 minutes to complete. Each CSES item is scored using a five-point Likert scale with response ranges from 1 (completely untrue) to 5 (completely true), with higher scores reflecting higher empathy levels. Previous psychometric testing of the CSES revealed good internal consistency with a Cronbach’s alpha (CA) of 0.96 for the overall scale.

### Data Analysis

Statistical analyses were performed using the statistical program SPSS (version 22). Data distribution was assessed by numerical methods included assessment of Z-score for skewness and kurtosis, Shapiro-Wilk’s test, and by visual methods including inspection of histograms and Q-Q plots for pre-test and post-test total scores. The Z-score is the number of standard deviations a value lies away from the mean and higher absolute Z-scores correspond to lower p-values. A Z-score of ±1.96 equates to a p-value of 0.05 in a two-tailed test. If data were normally distributed, parametric tests were performed, otherwise, non-parametric tests were used. The significance level was set at α < 0.05, two-sided.

### Comprehensive State Empathy Scale (CSEC)

To compare pre-test and post-test scores, the sum of the individual items of Comprehensive State Empathy Scale (CSES) for both the CSEC-Feelings and the CSEC-Perception were calculated. For both CSEC-Feelings scale and CSEC-Perception scale, due to small sample size in each study group and the fact that the majority of data did not comply with the assumptions of parametric tests (Shapiro-Wilk’s test, p<0.05), the non-parametric tests for data analysis were performed.

### CSEC-Feelings and CSEC-Perception

An individual Wilcoxon signed rank test was used to compare total calculated scores related to the CSEC-Feelings and the CSEC-Perception across the two measurement sessions (pre-test and post-test) in each study group by gender.

Additionally, a Kruskal-Wallis H test was conducted to evaluate if there were differences in the CSEC-Feelings scores and the CSEC-Perception scores across the three study groups in each measurement session (pre and post). Pairwise comparisons were also performed using Dunn’s procedure with a Bonferroni correction for multiple comparisons and adjusted p-values are presented. The distribution of the scores was also assessed by visual inspection of the boxplots. If data distribution were similarly shaped, medians were reported, otherwise mean ranks were used as a substitute score.

## Results

A total of 46 volunteers (36 female and 10 male) from three different undergraduate courses (nursing, landscape and architecture) participated in this study. Participants were aged from 19 to 37 years with an average of 23.8 years (SD= 3.65). While the majority of the female participants were from the nursing group (47.8%), most male participants were from the landscaping cohort (15.2%). The participants’ demographic and baseline characteristics are presented in Table 1.

**Table1.**
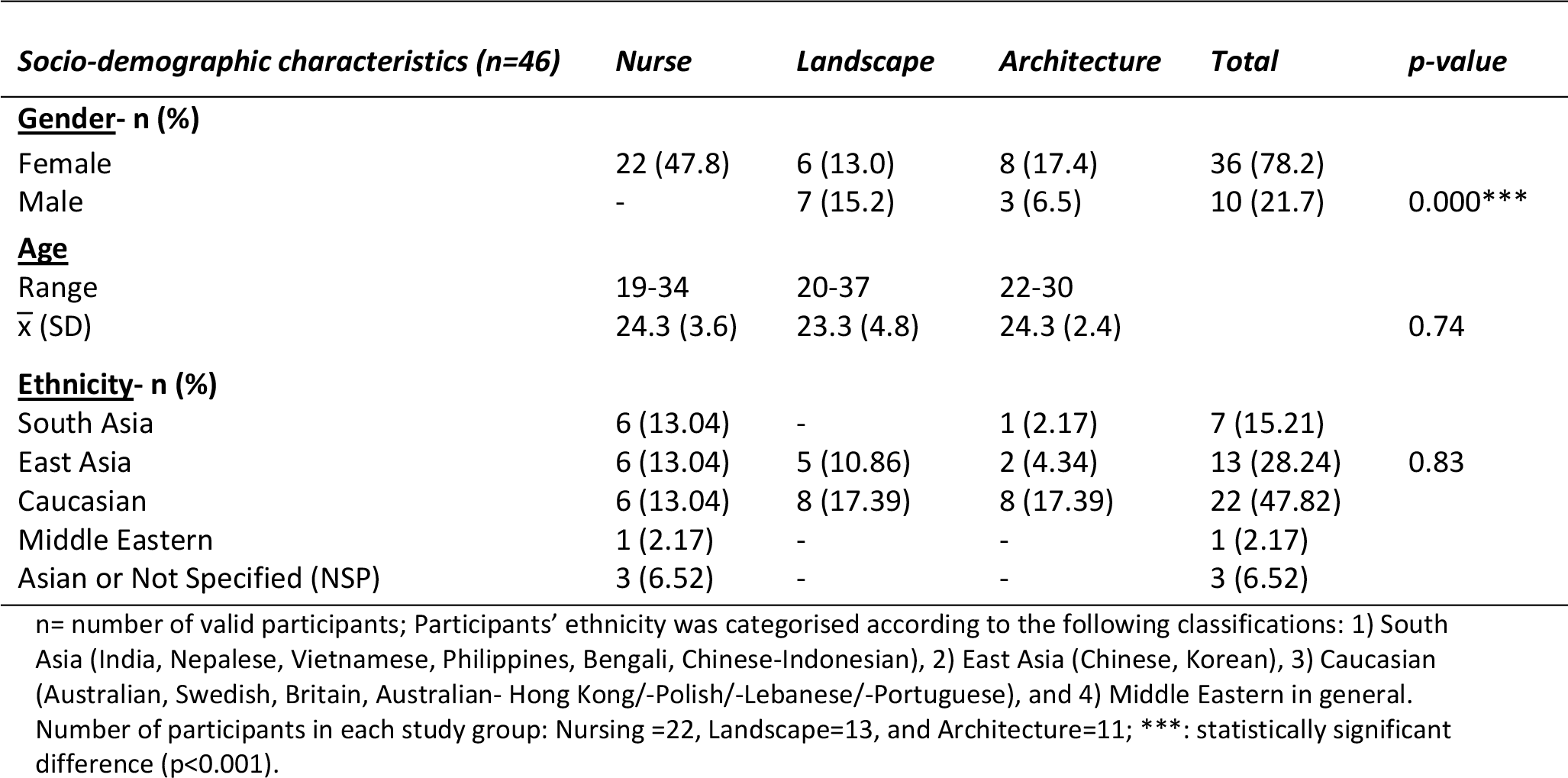
Socio-demographic characteristics of participants

Chi-Square test was used to compare gender and ethnicity differences among the three groups. However, when the expected frequency in any cell is less than five, then Fisher’s exact test is reported. There was significantly more female (n=36) than male (n=10) participants in the study (p<0.001). However, there were no significant differences in the ethnic categories across the three study groups (p=0.83).

The age difference between the three study groups was also analysed using ANOVA, and the result showed no significant difference in the mean age of participants among the study groups (p=0.74).

A Wilcoxon signed rank test was used to compare the pre and the post CSES-Feelings median scores in each study group by gender. For the positively worded items, the results showed a statistically significant difference between the pre and the post CSES-Feelings median scores only in the landscape group for both female and male participants but not for any other study groups, namely, nursing and architecture. Indeed, female participants in the landscape group reported a significantly higher level of CSES-Feelings when median scores from the pre-test (Mdn=12.5) were compared to the post-test (Mdn=19) (Z= −1.99, p<0.05). A similar pattern was also observed in the male landscape participants with an increased level of CSES-Feelings when the pre-test score (Mdn=14) was compared to the post-test score (Mdn=21) (Z=−2.20, p<0.05) (see Table 2).

**Table2.**
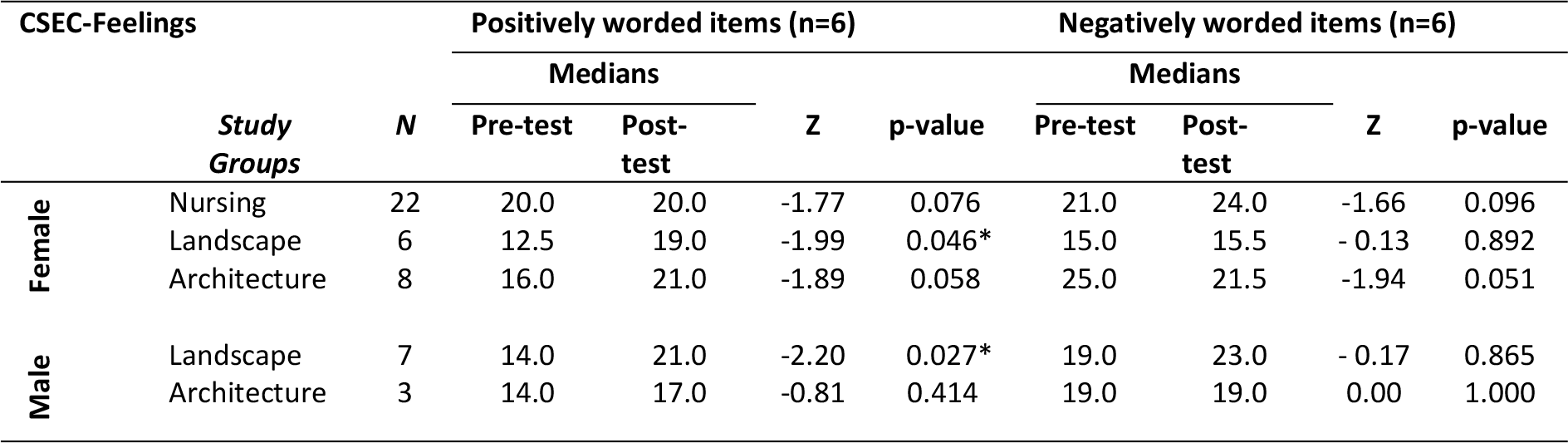
Pre and Post-test CSEC-Feelings comparison in each study group by gender. Comparison of CSEC Feelings’ level across the two-measurement time points (pre and post) within each study group; Female (n= 36) and Male (n=10); Wilcoxon Rank test (Z); Two-tailed significance level p<0.05.

A Kruskal-Wallis H test was conducted to evaluate the CSEC-Feelings scores across the three study groups in each measurement session (pre and post) by gender. The results showed that the distribution of CSES-Feelings scores were statistically significantly different between groups only for the positively worded items in the pre-test measurement session in the female group, *χ*2(2) = 12.288, p=0.002. The post-hoc analysis revealed statistically significant differences in CSES-Feelings scores only for positively worded items between the following study groups: nursing (mean rank=23.11) and landscape (mean rank=7.58) in the pre-test measurement (p=0.004) in the female participants (Figure 1).

**Figure. 1.**
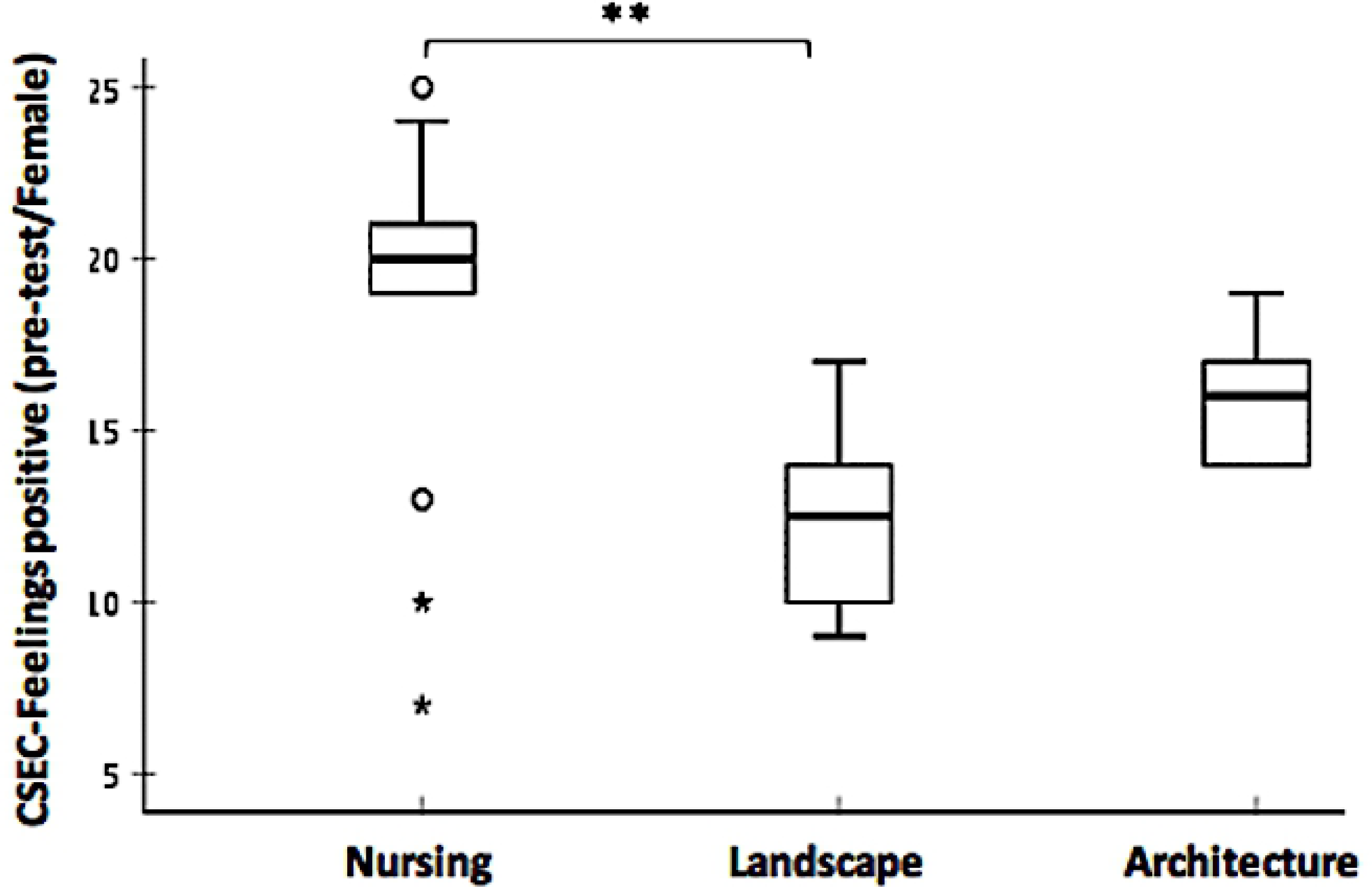
Comparison of CSES-Feelings median scores for positively worded items across the three study groups in the pre-test female group. The boxplots demonstrate comparison of CSES-Feelings median scores for positively worded items across the three study groups in pre-test (Nursing= 20, Landscape=12.5, Architecture=16) female group. The boxes are bound by the interquartile range (IQR) (top of the box represents the 75^th^ percentile, while the bottom of the box represents the 25th percentile). The boxes are divided by the median, and the whiskers attached to the box represent the minimum and maximum scores. Pairwise comparison demonstrated statistically significant differences in the CSES Empathy scores across different study groups (Kruskal-Wallis Rank Sum test, **p < 0.01; 2-tailed). Extreme values and outliers lied beyond the whiskers and denoted differently with a star and a circle respectively.

The participants’ CSES-Perception scores for each study group across the two measurement sessions were also evaluated by gender. Since the scores were not affected by gender, the data for the genders were pooled together. The results obtained from the analysis indicated no statistically significant difference in the CSES-Perception median scores across the two measurement sessions in each study group. Descriptively, however, the CSES-Perception median scores related to both the landscape group and the architecture group presented a slight increase, as opposed to the nursing group wherein there was a slight decrease in the CSES-Perception median scores when data related to the pre-test compared to the post-test (Figure 2).

**Figure 2.**
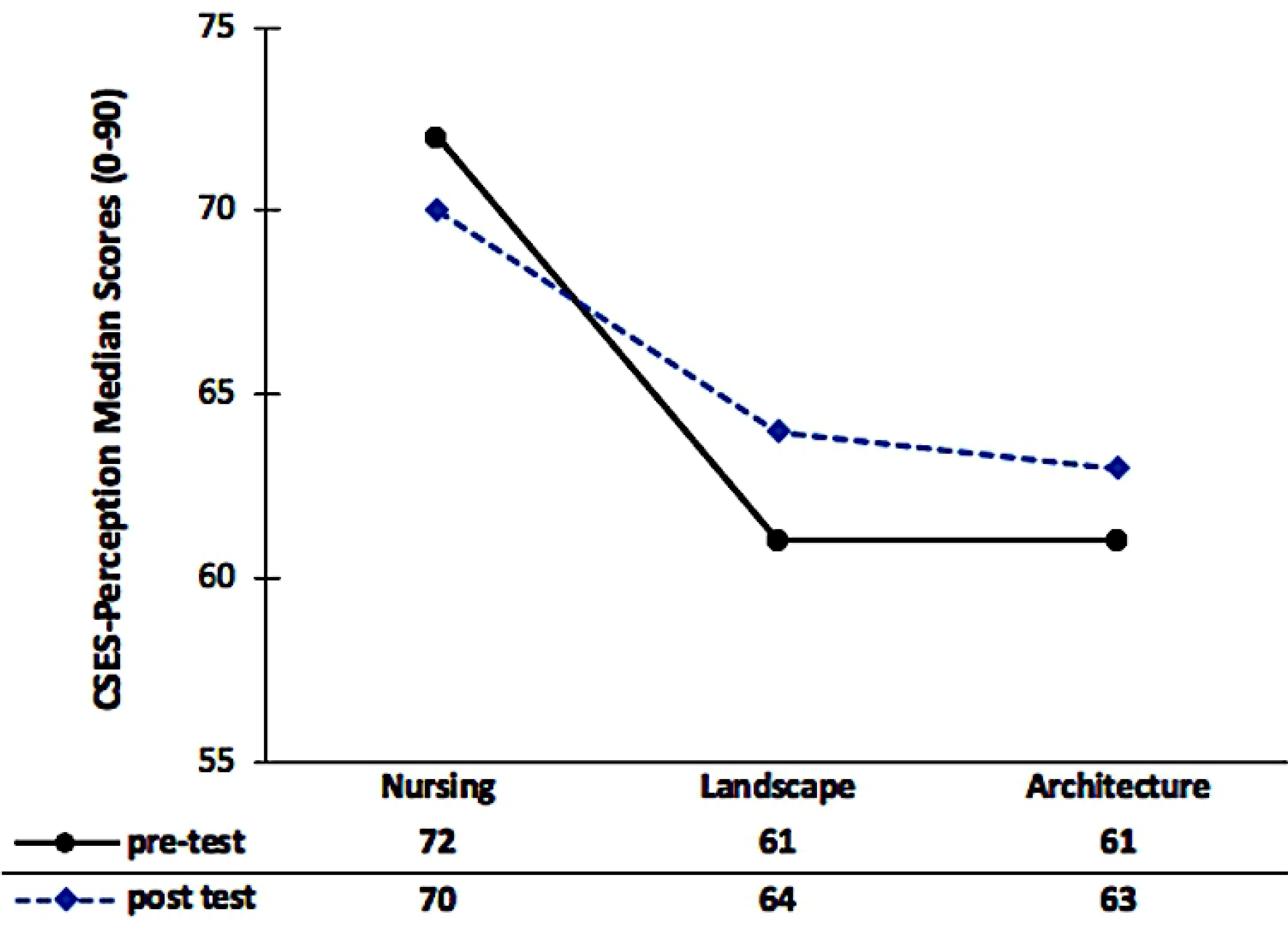
CSES-Perception median scores in each measurement session across the three study groups (nursing, landscape, architecture)

Furthermore, a Kruskal-Wallis H test was conducted to evaluate the CSEC-Perception across the three study groups in each measurement session (pre and post) by gender. The distribution of CSES-Perception scores were statistically significantly different between study groups for both the pre-test measurement session (*χ*2(2) = 9.530, p=0.009) and the post-test measurement session (*χ*2(2) = 8.039, p=0.018) only in the female group. The post-hoc analysis revealed statistically significant differences in CSES-Perception scores between the following study groups: nursing (mean rank=22.36) and landscape (mean rank=11.00) (p= 0.05), and nursing and architecture (mean rank=11.81) (p=0.040) in the pre-test measurement in the female participants (Figure 3A). However, the post-hoc analysis in the post-test measurement session was only a statistically significant difference in CSES-Perception scores between nursing (mean rank=21.98) and architecture (mean rank=11.31) (p=0.036) (Figure 3B).

**Figure. 3A-3B.**
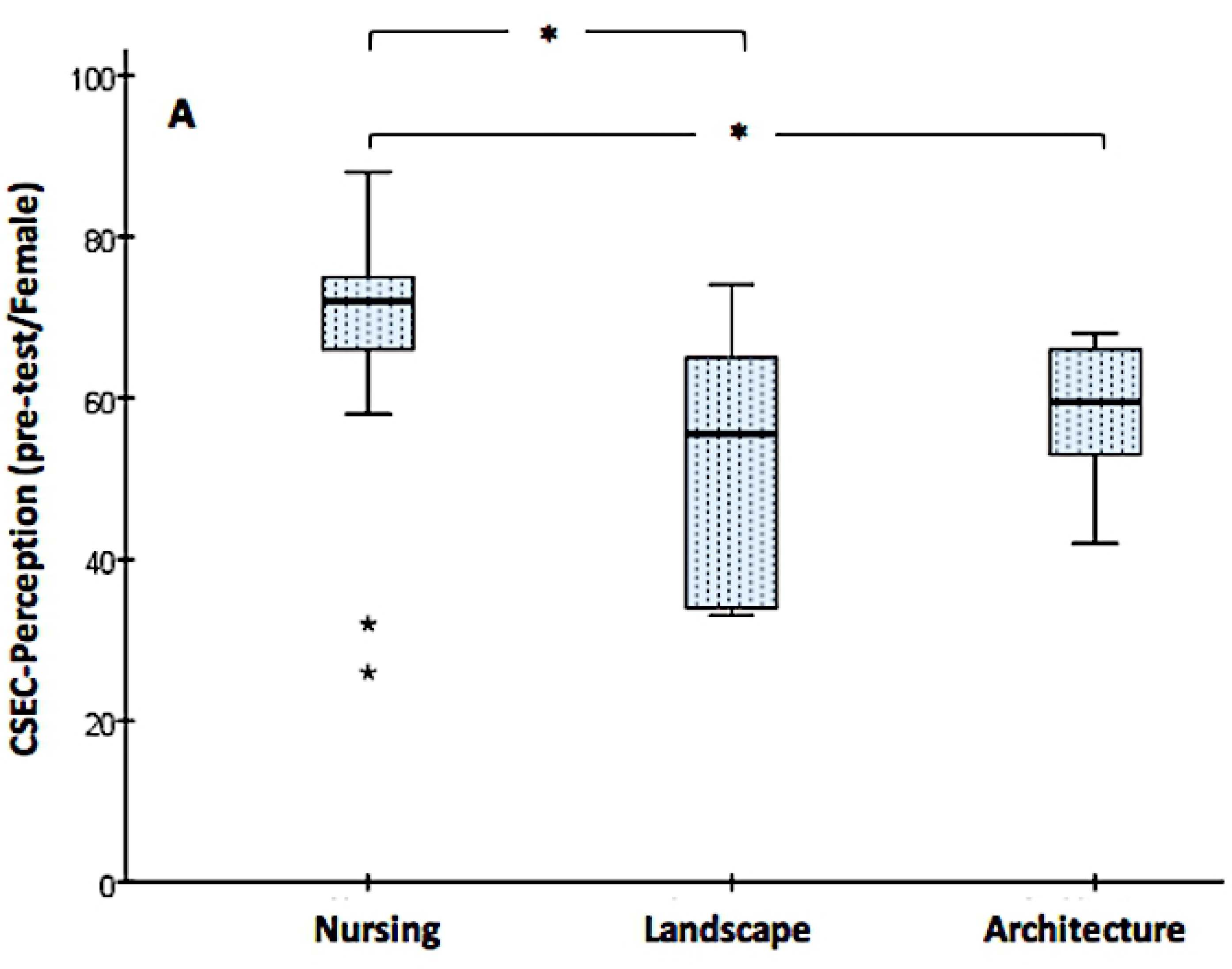
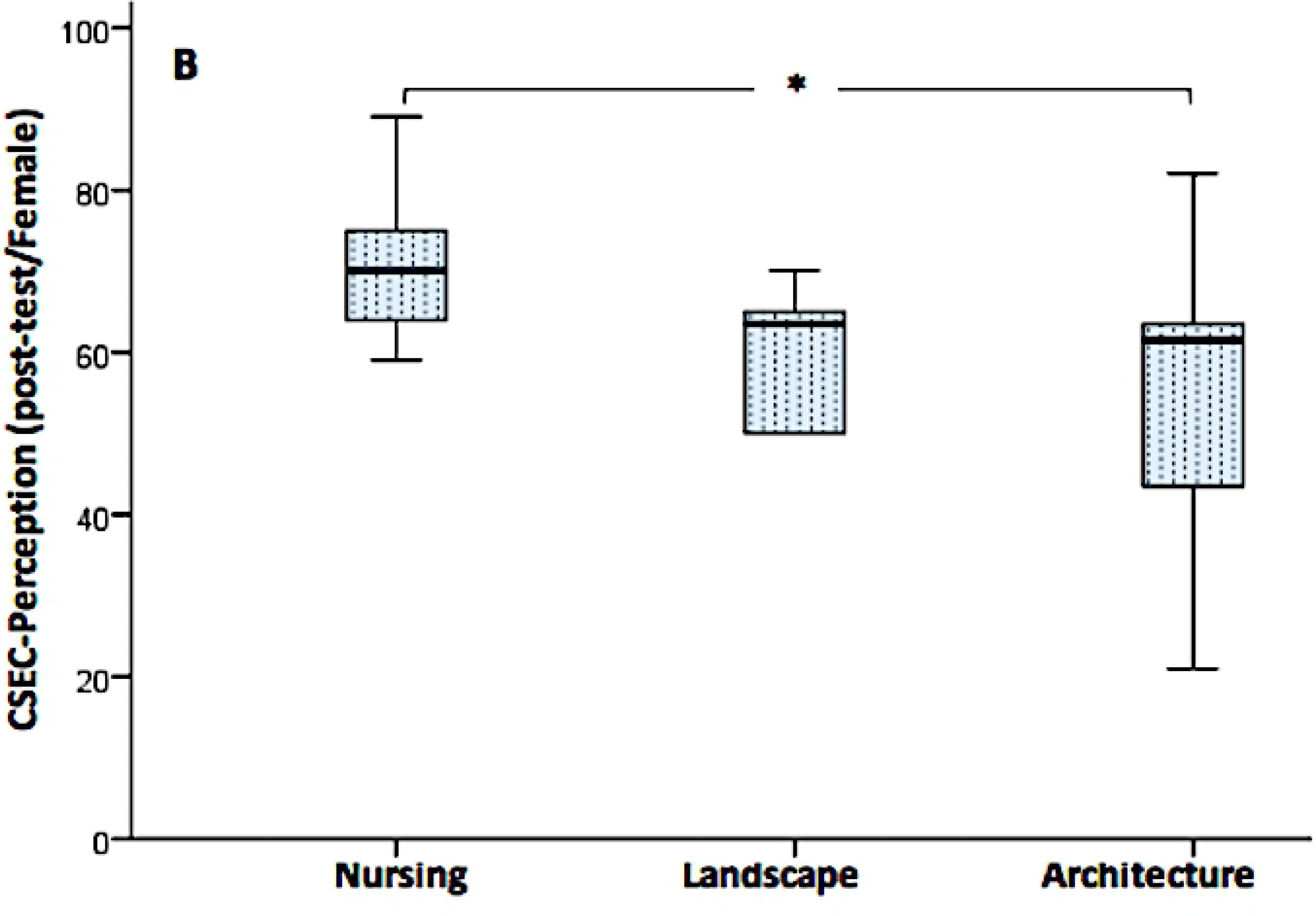
Comparison of CSES-Perception median scores across the three study groups in both measurement sessions (pre-test and post-test) in the female group. The boxplots demonstrate comparison of CSES-**Perception** median scores across the three study groups in the pre-test, 3A (Nursing= 72, Landscape=56, Architecture=60) and the post-test, 3B (Nursing= 70, Landscape=64, Architecture=62). The boxes are bound by the interquartile range (IQR) (top of the box represents the 75^th^ percentile, while the bottom of the box represents the 25th percentile). The boxes are divided by the median, and the whiskers attached to the box represent the minimum and maximum scores. Pairwise comparison demonstrated statistically significant differences in the CSES Empathy scores across different study groups (Kruskal-Wallis Rank Sum test, *p < 0.05; 2-tailed).

## Discussion

In Australia, one in six women have experienced physical or sexual violence by a current or former partner [26], yet reports suggest that many people are judgemental and fail to consider the complexity of social issues related to domestic violence [27–28]. Domestic violence is the single largest cause of homelessness for women [1] and the greatest health risk factor for women aged 25-44 [29]. Homelessness can also result in discrimination which restricts people’s access to necessary services and supports, including housing and employment. [30] However, empathy is one of the strongest negative correlates of prejudice [31] providing a lens through which women experiencing homelessness can be better understood and addressed.

The study described in this paper sought to examine impact of an authentic interdisciplinary learning experience on nursing, architecture and landscape students’ empathy towards women and children who access refuge services as a result of homelessness and/or domestic violence. Prior to the study we assumed that there would be a pre-post increase in empathy levels in all students, but that the increase may be less for nursing students who were frequently exposed to situations and scenarios designed to increase empathy towards vulnerable groups.

The results indicated that nursing students’ pre-test empathy levels were markedly higher than those of architecture and landscape students, and that changes in pre-post empathy scores for this cohort was minimal. This possibly reflects a dispositional tendency – nursing fundamentally involves a commitment to understanding and responding to human suffering – so that the nursing students commenced the study with higher levels of empathy and were therefore less likely to experience a significant change over the 12-week study period.

The increase in landscape students’ empathy was approximately twice as much as the nursing students, whereas architecture students’ empathy levels decreased by a similar amount. These results were unanticipated outcomes of the study but may be explained, at least in part, by the gender and age distribution of the participants.

There was significant variation in gender within the overall sample, with more male than female landscape students, compared to architecture students who were predominantly male and nursing students who were all female. The generally accepted fact that males are less empathetic than women [32] may have influenced by the gendered nature of the results. The range in ages, and the relationship between life experiences and empathy levels may have also influenced the results as the landscape students were a slightly older group, although this is only an assumption. Additionally, no clear relationship could be found between empathy levels and ethnic diversity, with the sample that was predominantly South East Asian and Caucasian, and a small number of Middle Eastern students.

Other reasons for these results include the nursing students’ greater exposure to refuge users in their field of study, and consequently, their understanding of human suffering may be more refined. Architecture and landscape students may have never dealt with design in the context of social issues which include human cruelty and suffering. The teaching and learning activities for the three cohorts of students, each enrolled in different subjects, may also have contributed to the outcomes. For example, architecture and landscape students worked from a data set provided by conversations with stakeholders and independent research, with input from nursing students – to produce a design proposal. This hypothetical project was presented for evaluation by a select number of academics with little expertise in the field of domestic violence. Design students, aware of this mode of assessment, may have focused their design responses on aesthetic concerns in an aspirational effort to appeal to a broader audience of judges. Nursing students, in contrast, may have responded more directly to the needs of women accessing a refuge with learning activities that involved a more empathetic framework to be established. Additionally, the nursing students undertook a four-week work-based placement in various women’s health service environments, while the architecture students focused on the typology of refuge, exploring retreats, monasteries, sanatoriums and domestic spaces as precedents, and landscape students developed a scheme for a real-life refuge, which may also have influenced the outcomes of the study.

The CSES levels of architecture students at the pre-test stage might also be explained in part by response bias theory which suggests that experimental conditions can bias respondents thereby damaging the validity of a study. Architecture and landscape students are used to rigorous competition to perform and present unique and critically challenging ideas. The first responses were higher - when the desire to ‘please’ the instructor was greater, than at the post-test stage when the students might have deduced that the study had little or no impact on their results [33]. Architecture students may be positively distracted by the requirement for social agency in the outset of the course, but in the finalising of the subject deliverables revert to practices of ensuring aesthetic quality is privileged, therefore removing emphasis from the importance of the user.

### Limitations

The results of this study are limited by the relatively small sample size and the fact that the participants were from one university which prevents generalisability and representativeness. Thus, the results should be interpreted with caution and indicate the need for ongoing research. Additionally, limitations inherent to the two-group pre-test post-test research design such as the lack of control group and reactive interaction effect of pre-testing must be considered when evaluating the internal and external validity of the study. Lastly, the outcomes from this study demonstrated an immediate change in participant empathy scores only. Whether these results would be sustained over time and, more importantly, whether they would influence participants’ professional practice, has not been determined.

The potential areas of this project which still remain unresolved include the specificity of the CSES in interdisciplinary subjects and whether it can be effectively used to measure radically different disciplines equally. The impact of response bias on students, particularly of a design background, is still not completely understood. Lastly, the development of empathetic understanding in different genders of students has not been clearly tracked in this project but may in fact have a large effect on the resulting data.

### Future research

Future studies should specifically document each collaborative session and its impact on empathy levels to decipher which activities are more effective in enhancing students’ empathy levels. Future iterations of this project could also include a wider range of discipline specific courses, such as law, building, engineering and business.

This project assisted students to develop the requisite skills of empathetic understanding while still in a university setting rather than relying on this development to occur in the future practice of the professional. In providing more of these opportunities for students during their university studies, their ability to further develop skills in empathetic understanding and user-based responses may be enhanced.

## Conclusion

Through providing students with an authentic learning experience which has obvious social impact and currency, students were enabled to reflect on their existing perceptions and to gain new insights into the concerns, preferences, and perspectives of individuals who access and provide refuge services. This teaching and learning strategy had the potential to enhance students’ understanding of service user requirements so as to better design safe spaces that are conducive to the needs of women and children who have experienced domestic violence. This project provides a model for future variations of this interdisciplinary subject as well as for other social impact projects within a university context.

## Acknowledgements

The authors would like to acknowledge the support of the following: UTS Social Impact Research group; Board Director and staff at Hornsby Ku-ring-gai Women’s Shelter; Mayor of Ku-ring-gai Council; UTS Centre for Social Justice and Inclusion, students who participated in these subjects and the schools within which the subjects were facilitated.

